# Systemic Commitments of the Context-Dependency; Basics of Elementary Coordination in Circadian Cycles

**DOI:** 10.1101/738872

**Authors:** Chulwook Park

## Abstract

The present study was attempted to measure whether the dynamics of elementary coordination is influenced by an overarching temporal structure that is embedded in circadian rhythms (part 1) as well as the systemic proof associated with the intelligent capabilities (part 2). For part 1, evidence of entrainment or any influence of the embedding rhythm were examined on the stability or attractor location. The estimations from the dynamics of the relative phase between the two oscillations show that while (i) circadian effects under the artificially perturbed manipulation were not straightforward along the day-night temperature cycle, (ii) the circadian effect divided by the ordinary circadian seems to be constant along the day-night cycle. Corresponding to this evidence related to performance consequences depending on the organism and environmental interaction, the part 2 determined the impact of circadian (mis)alignment on biological functions and raised the possibility that the disruption of circadian systems may contribute to physical complications. The observations entail rules that self-attunement of current performance may develop not at a single component but across many nested, inter-connected scales. These inter-dependencies from different object phase may allow a potential context-dependent explanation for goal-oriented movements and the emergent assumption of a principle of organisms embedded into their environmental contexts.

---

Men, earthworms, single-cell organisms, and even plants are aware of their surroundings and organize actions according to their circumstances (Cariani, 1993; Ford, 2008; Strong & Ray, 1975). Exhibits of autonomy and control of functions have been the goal in contemporary research to explain agency more scientifically (Shaw & Kinsella-Shaw, 1988). To understand this directed behavior as a call for the organism-environment system rather than as merely pertaining to the organism (Turvey & Carello, 2012), we seek to expose the laws that underlie intelligent capabilities.

One useful strategy looks for cycles at all time scales and aims to show how interacting cyclic processes account for the emergence of new entities, many of which are similarly cyclic (Iberall, 2001). The central idea is that the earth’s cycles—the geophysical, hydrological, meteorological, geochemical, and biochemical—have interacted to create self-replicating living systems abiding particular cyclicities (Soodak & Iberall, 1978). We inquired into whether something akin to attunement to the environmental 24 day/night temperature cycles (Maury, Ramsey, Bass, 2010) may be apparent in an experimental setting of elemetnary coordination, a setting that has been used to examine self-organization in biological systems (Kugler & Turvey, 1987). We asked whether unintentional coordination with an environmental rhythm might raise by assessing whether the dynamics of bimanual coordination is influenced by an overarching temporal structure that is irrelevant to the task (see Part 1).

We should embrace, or at least acknowledge, methodological and theoretical diversity given any observations. As one of the prototypes, circadian systems can be considered to be a comparatively simple frame for dealing with complicated physical phenomena. Presenting strategy that includes very many processes and parameters qualitative understanding (Stepp & Turvey, 2009). We take it that to be proof for the possibility as a way of mechanistic principle in living systems in order to address its pervasiveness. This makes us to its mechanistic identifiable in that how components of certain structures or elementary interact dynamically for that phenomena (Bechtel, 2009a,b). The circadian oscillation considered must be incorporated as an endogenous system that represents of the primary clock in both biology and environment, it is worthwhile to be contacted with appropriate methodological basic (Stepp, Chemero & Turvey, 2011). Contributing from this assumption may open a hypothesis of coupling of organism and environment, as an alternative to understanding systems dynamics (Warren, 2006) (see Part 2).

## Part 1: Embedded Motor Synchrony in Circadian Rhythm

The present study of this section does so in two principal ways in order to determine the physical characteristics in an effort to find that approximations under certain conditions serve these self-potentials. The first involves an increase in the capability to self-generate forces along the lines of the roles of the fundamental dimensions of environments (temperature embedding in light-dark cycles). The second tied to observe availability of an internally based source (coordination) or sources of force (stability and entropy) within dynamical boundaries in systematic ways. We develop a general model for the experimental designs to obtain any influence of the embedding rhythm as follows.

### Model of the experimental design

Formation and retention refer to propriospecific information about the states of the muscular-articular links, and the dynamical criteria of the stability pattern constrain the patterns or characteristics. To be specific, let us consider a qualitative physical system such as stiffness, damping, and position over time in a dynamical mass-spring system as given.

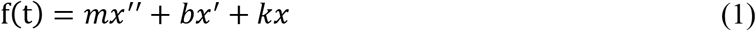

Here, *m* is mass, *b* is friction, and *k* denotes the stiffness. The variable t is time, χ denotes the position, χ′ is velocity, and χ″ represents acceleration. In physics, because damping is produced by a process that dissipates the energy stored in the oscillations, the interplay between input and damping approaches a stationary fixed point in the long-time limit.

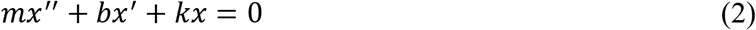

Such systems possess a static equilibrium point, which is called a point attractor (Kugler, Kelso, & Turvey, 1980). The property of this dynamic has been applied not only to a physical system but also to descriptions of the human neuromuscular level (Kugler & Turvey, 1987). This function involves an investigation of the intact movement of a limb oscillator in terms of muscle-joint kinematic variations (kinematic position, velocity, acceleration) over time. When we are asked to swing two limbs comfortably, this can be characterized by the pendulum’s dimension (Pikovsky, Rosenblum, & Kurths, 2003), namely, simplifying the point attractor while restricting it to certain domains of phase space [(*θ2 – θ1* ≈ 0), (02 – 01 ≈ *π*)]. In this equation, with the phase difference, 02 – 01 ≈ 0 denotes a condition of nearly synchronized in-phase, and *θ*2 – *θ*1 ≈ *π* indicates that this in an anti-phase. The observed relative phase or phase relation (ϕ) between two oscillators at ϕ ≈ 0 deg (inphase), or ϕ ≈ 180 deg (anti-phase) have been modeled as the point attractors in our limb system, as they are purely stable patterns (Turvey, 1990).

### Elementary coordination

In the observed relative rhythmic segments patterns, the in-phase ϕ = 0 condition is more stable than the anti-phase ϕ = π condition. Inspired by a number of studies on the 1:1 frequency locking of the left- and right-hand phase defined as ϕ = (*θ_L_* – *θ_R_*)—the difference between the left (L) and right (R) phase angles (ϕ)—has led to the identification of important invariant human system features (Kelso, 1984).

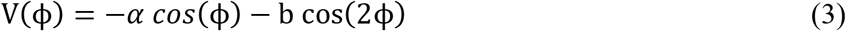

In this equation, ϕ is the phase angle of the individual oscillator. In addition, α and b are coefficients that denote the strength of the coupling between the two oscillators. The relevant regions of the parameter space allow the potential V(ϕ); the negative signs in front of the coefficients simplify the equation of motion. A relative 1:1 frequency-locked coordination phase [V(ϕ)] is determined by the differences between the continuous phase angle [–*α cos*(ϕ) – b cos(2ϕ)] of the oscillator’s two components: the stability of the point attractor can be varied by varying the pendulum’s dimensions.

This function indicates that the minima of the potential are located at ϕ=0, and that ϕ = ±π (Haken, Kelso, & Bunz, 1985). Given this scenario, the function can be estimated in terms of how the potential will change in shape, as the control parameter (energy cost) increases. Based on the observed mechanism for the point attractor with a simple function, the present model proposes the in-phase bimanual rhythmic coordination synchrony pattern as a particularly well-suited physical source. This allows a useful reference for system stability coordination tasks in which this functional pattern can be applied to all human movement, muscles, and even a neural network. Actual intersegmental coordination, however, is additionally shaped by the contingencies of adjusting to environmental vagaries. How these extrospecific requirements and information types are incorporated into the physical stability patterns can be assumed by the level of symmetry coordination (Amazeen, Amazeen, & Turvey, 1998). In order to harmonize the effects of motor stability toward environmental symmetry, this study investigates the following elaboration.

### Symmetry breaking in bimanual coordination dynamics

The potential [V(ϕ)] extends the described assumption in terms of the difference between the uncoupled frequencies of bimanual rhythmic components [Δ*ω* = (*ω_L_ — ω_R_*)]. Where *ω* is the preferred movement frequency of the left (*ω_L_*), right (*ω_R_*) individual. If the relative phase between *ω_L_* and *ω_R_* were equal (Δ*ω* = 0), this pattern would be assumed to be a perfectly identical symmetry. However, the preferred movement frequencies of the individual oscillators in in-phase are large (i.e., function: b/a=0.5, detuning=-0.5, or detuning=-1.5), the expected stability of the rhythmical limb oscillation dynamics become greater than equal.

Such phenomena of the symmetry breaking must be another fundamental feature of the coordinative system (Amazeen et al., 1998). From this dynamic, a different noise of the underlying subsystems (neural, muscular, and vascular) can be estimated around an equilibrium point, and this might conceptualize the model when it comes to making operational definitions of each category in which the model has to consider the variability of the relative phase frequencies between two limbs:

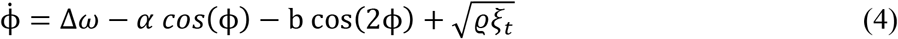

The estimation of two oscillators’ relative phase 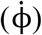 is captured by the parameter (Δ*ω*) of the preferred movement frequency of the individual segment [*α cos*(ϕ) — b cos(2ϕ)] with the noise 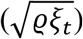. Given the equation of the preceding model (grouped as the kinematics of motor stability according to the coordination task of synchronization), such a term has been used to capture purely functional dynamics regarding the equilibria and is confirmed usually as in the time and temporal difference between an oscillating limb (Treffner & Turvey, 1995).

Researchers (Treffner & Turvey, 1995), conducting experiments in handedness, advanced the elementary coordination dynamics. They added two add (sine) terms for the coefficients, whose signs and magnitudes determine the degree and direction of asymmetry, as follows;

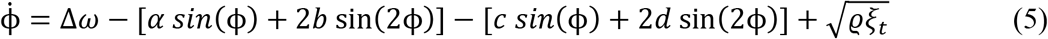

Here, 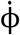 indicates a coordination change. Δ*ω* refers to a symmetry breaking through frequency competition between two limbs. [*α sin*(ϕ) + 2*b* sin(2ϕ)] denotes a symmetric coupling defined by relative phase of 0 and π attractors (this form of the term could be derived as the negative gradient potential V with respect to ϕ); and the [*c sin*(ϕ) + 2*d* sin(2ϕ)] terms means added asymmetric coupling attractors with the stochastic noise 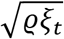. This extended equation refers to the fact that the emergent elementary dynamics between limbs or limb segments was governed by a slightly asymmetric potential of the [*c sin*(ϕ) + 2*d* sin(2ϕ)]. That suggests extended collective dynamics of the inter-segmental rhythmic coordination of the periodic components.

### Thermoregulatory symmetry breaking of the elementary coordination

Inspired by the complementary symmetric and asymmetric influences, the described model was applied to investigate the difference between the coupled or uncoupled frequencies of the temperature-rhythmic components between the core body and circadian cycles.

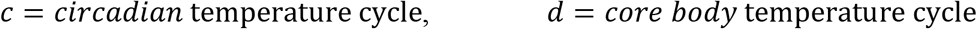

Where *d* is the preferred rhythmic frequency of one (the homeostasis cycle) and another (*c* = circadian cycle) individual. Whereas b/a determines the relative strengths of the fundamental in-phase equilibria, small values of *c* and *d* break the symmetry of the elementary coordination dynamics while leaving their essential coupling characteristics.

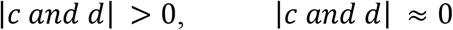

In this proposed assumption, the coefficient of the *d* should be more important, producing the empirically observed perturbation in the equilibrium phase state, and then the *c* should be set to zero without loss of generality, given that we cannot manipulate the environmental circadian cycle. As one can see, if the coupling between *d* and *c* is strong (|*c and d*| ≈ 0), this pattern would potentially be expected to be in perfectly corresponding symmetry with the environmental requirement. However, the preferred rhythmic coupling of individual oscillators in an in-phase condition becomes a difference (|*c and d*| >0), and thus the expected stability or variability of the rhythmical-component oscillation dynamics will become greater than equal. Given the preceding assumption (grouped as the kinematics of motor stability according to the coordination task of synchronization), the equation was extended to a novel task in which there are different sources of symmetry breaking through thermal variables, as information has not yet been made available about the effects of bimanual dynamics in instruction on circadian temperatures.

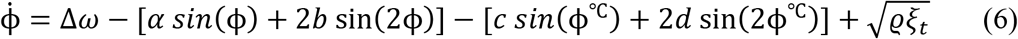

In this equation, in the bimanual 1:1 rhythmic coordination performed at different coupled frequencies, the symmetric coupling coefficients will be not the same. There will be an increase in detuning (Δ*ω*) and a decrease in the relative strengths of the attractors at 0 and *π*. However, when it comes to our limiting case of Δ*ω* = 0 on the approximately identical symmetry temperature parameters (core body and circadian cycle), what should we expect? The final estimation between the relative phases of two oscillators 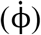 will be captured mainly by the parameter of the asymmetric thermoregulatory coupling [*c sin*(ϕ°^C^) + 2*d* sin(2ϕ°^c^)] with noise 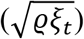. From this dynamic, the different noise types of the underlying subsystems (neural, muscular, and vascular) around an equilibrium point were able to be estimated, suggesting that such phenomena related to symmetry breaking may be yet another remarkable feature of the coordinative system.

In sum, this experiment was required to have a condition of in-phase (ϕ = 0) oscillated simultaneously at the 1;1 frequency locking. The same goal using the functional symmetry dynamics of different effectors will be influenced by the asymmetric thermal regulation symmetry breaking through both circadian temperature cycles. Namely, the effect of one of the contralateral homologous relative limbs phase might be not identical to the impact of the others. The expected stability pattern, from intuition given a different motor, appears to allow the biological symmetry dynamic to be understood in the ecological context. This attunement to the circadian temperature approach implies an emergent property of the system.

### Method of the first experimental design

Experiments 1 embedded a bimanual coordination task in an ordinary 24-hour day-night cycle (5:00, 12:00, 17:00, and 24:00). Data sets were subjected to an analysis of variance. The setting asks “Is our system influenced by an ecological feature?” A metronome was used to impose a rhythm reflecting the natural period of the pendulum system. The ordinary temperature condition served as a replication of standard in-phase bimanual coordination tasks. The participant was seated with his or her arms supported on armrests and a pendulum in each hand held firmly to prohibit within-grasp motions. Gaze was elevated to prevent viewing the pendulum oscillations which arose from motions strictly about the wrist. Experiment 1 (n = 8) used a setting that minimized variability: in-phase coordination of the two pendulums (relative phase *ϕ* = 0°) with no detuning (i.e., the two pendulums had the same eigenfrequency). In the experiments, bimanual coordination was performed at a metronome beat (1.21 s) — this period was chosen because it corresponded to the natural period of the pendulum system: Amazeen, Amazeen, Trffner, & Turvey, 1997—without concern over amplitude or frequency (see Table 1).

**Table 1:** Data collection for the experiment 1. 8 participants, 4 circadian points, 6 trials at each circadian point. Note: Participants were asked to swing in-phase of their limbs in different anatomy points [192 data set (3-level: wrist, elbow, and shoulder)], but used only wrist joint data (64 set) for analysis. Duration of each trial is 1 minute and 5 minutes rest interval between trials.

| Condition | Participants (N) | Circadian points | Trials at each circadian | Task/rest (min) |
| --- | --- | --- | --- | --- |
| Normal | 8 | 5:00 | 6 | 1/5 |
|  |  | 12:00 |  |  |
|  |  | 17:00 |  |  |
|  |  | 00:00 |  |  |

### Results of the first experimental design

A variety of measures (e.g., phase shift, variability, entropy) were examined for evidence of entrainment or any influence of the embedding rhythm on stability or attractor location. Absolute differences in parameters (fixed point shift, variability as a function of frequency competition, and entropy production) were found, especially, at circadian 17:00 point. The parameters changed in the opposite direction from the core body temperature. While the core body temperature rhythm shows a minimum at 05:00 h but has a maximum at about 17:00 h, behavioural performance (entropy; Shannon, 1948) shows a maximum at 05:00 h but has a more clearly defined minimum at about 17:00 h in day-night cycle (see Figure 1).

**Figure 1.**
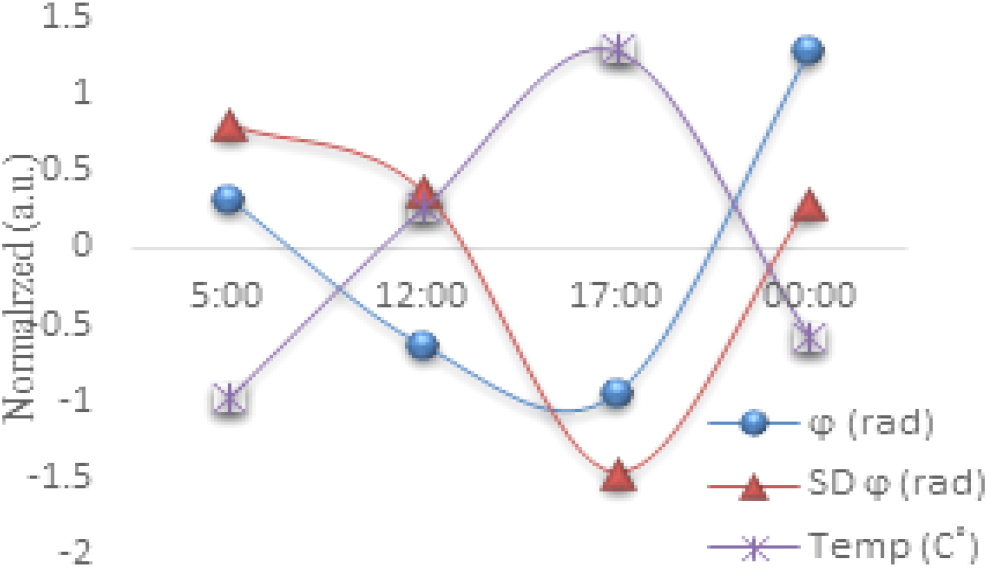
General tendencies in the normal condition. Normalized = standard score (Z calculation), a.u. = arbitrary unit, blue line = fixed point shift, red line = variability as a function of the frequency competition, Temp = temperature (Celsius), 5:00, 12:00, 17:00, and 00:00.

The result shows the general feature of the average trend in ordinary circadian cycles. As shown in the table, the main effect of the uncertainty on the ongoing circadian cycle was not significant [*F*(1, 3) = 1.074, *η*^2^ = .823, (*p* < 0.376)]. Absolute differences in the widths of the circadian cycle between temperature and entropy production can be observed, especially at the circadian points of 5:00 and 17:00 (t = 1.764, *p* < 0.103).

### Method of the second experimental design

Experiments 2 and 3 asked, “How does our system adapt to regular or irregular thermal structures?” Normal and abnormal day-night circadian temperature effects were compared at dawn (5 a.m., approximately when core temperature reaches its minimum) and dusk (5 p.m., approximately when core temperature reaches its maximum; Aschoff, 1983). In-phase coordination without detuning was performed at dawn and dusk. A metronome was used to impose a rhythm reflecting the natural period of the pendulum system. In addition, a short-term, thermodynamic manipulation was introduced. Prior to half of the sessions, participants (n = 8) donned a heated vest for 30 min that increased their core temperature by certain (within 0.5°, *SD* 0.2°) degrees (Exp 2). And participants (n = 8) donned an ice vest for 30 min that reduced their core temperature by certain (within 0.4°, *SD* 0.15°) degrees (Exp 3). The expectation was that the thermal (increasing or decreasing) manipulation’s influence on coordination would interact with a time of day (see Table 3, 4).

**Table 2:** Experimental result 1. Each type of raw value for the normal day-night temperature effects. P denotes the participants, numbered 1-8, *υ*_ave_ fixed point shift, *SD_υ_* variability as a function of the frequency competition. T = core body temperature (Celsius).

|  | Circadian 5:00 |  | Circadian 12:00 |  | Circadian 17:00 |  | Circadian 00:00 |  |
| --- | --- | --- | --- | --- | --- | --- | --- | --- |
| | $\phi_{ave}$ | $SD\phi$ | $\phi_{ave}$ | $SD\phi$ | $\phi_{ave}$ | $SD\phi$ | $\phi_{ave}$ | $SD\phi$ |
| P1 | 1.1 | 6.0 | 0.8 | 5.2 | 1.5 | 6.7 | 6.0 | 43.1 |
| P2 | 4.0 | 43.1 | 2.8 | 10.8 | 5.0 | 6.2 | 3.8 | 16.0 |
| P3 | 11.2 | 6.7 | 4.0 | 6.0 | 1.5 | 5.4 | 3.9 | 7.7 |
| P4 | 2.67 | 77.3 | 1.7 | 83.7 | 6.6 | 10.1 | 7.6 | 6.7 |
| P5 | 6.18 | 40.7 | 3.0 | 13.6 | 2.7 | 11.3 | 4.6 | 20.6 |
| P6 | 1.95 | 38.0 | 5.6 | 59.7 | 5.8 | 33.3 | 4.62 | 41.1 |
| P7 | 3.52 | 40.1 | 4.4 | 20.9 | 0.8 | 6.1 | 3.8 | 55.2 |
| P8 | 4.25 | 17.3 | 5.5 | 40.0 | 1.9 | 35.9 | 7.1 | 44.1 |
| T(C°) | 36.6 |  | 36.8 |  | 37.0 |  | 36.6 |  |

**Table 3:** Data collection for the experiment 2. 2 conditions, 8 participants, 2 circadian points, 6 trials at each circadian point. Note: Participants were asked to swing in-phase of their limbs in different anatomy points [192 data set (3-level: wrist, elbow, and shoulder)], but used only wrist joint data (64 set) for analysis. Duration of each trial is 1 minute and 5 minutes rest interval between trials.

| Condition | Participants (N) | Circadian points | Trials at each circadian | Task/rest (min) |
| --- | --- | --- | --- | --- |
| Normal | 8 | 5:00 | 6 | 1/5 |
|  |  | 17:00 |  |  |
| Abnormal<br>(heat based) | 8 | 5:00 | 6 | 1/5 |
|  |  | 17:00 |  |  |

### Results of the second experimental design

Systems state was estimated from the dynamics of the relative phase between the two limbs oscillating at the wrists. Stability of the performance was affected by the temporal locus during the circadian cycle, as well as the introduction of the heated vest (Exp 2) and the ice vest (Exp 3); the influence of the thermal manipulation was not identical. Even if the same external temperature perturbations were given, the influence of the vest was negatively exaggerated (increasing entropy) at dawn, but the influence of the vest was positively exaggerated (decreasing entropy) in the evening (see Figure 3, 4).

**Figure 2.**
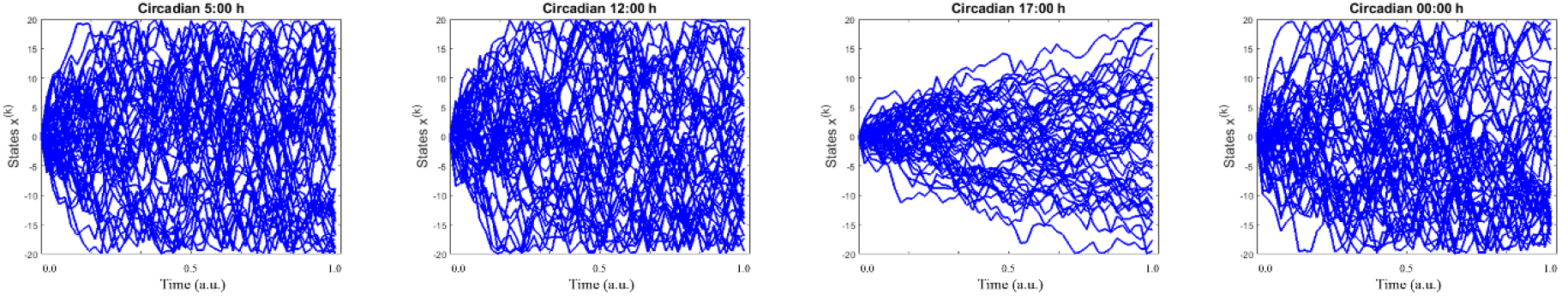
Uncertainty characteristics in the normal condition. The figures denote the estimated entropy states according to the time series. Left = 5:00, middle left = 12:00, middle right 17:00, right = 00:00. Note: for this realization, participant 3’s data was used which could be represented as the most similar scores with the arbitrarily normalized scores from the averaged all participants’ scores as a function of the frequency competition [*SDϕ* (rad)].

**Figure 3.**
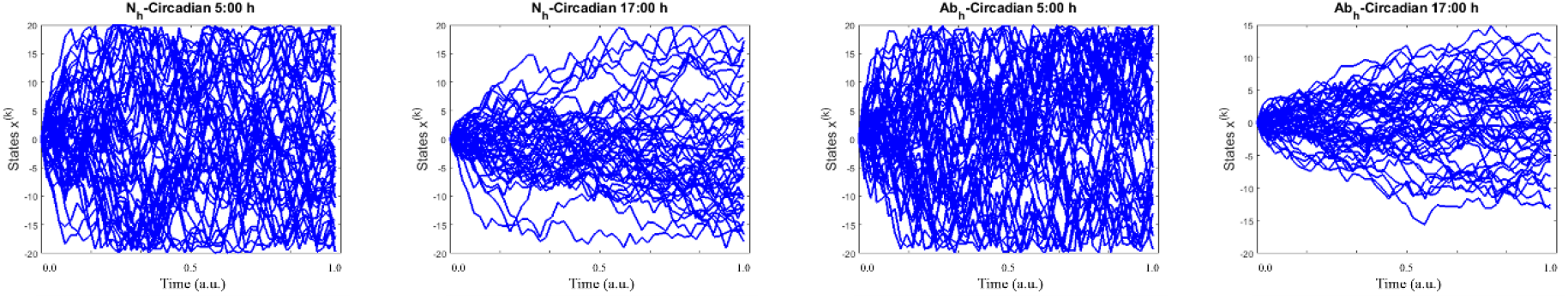
Uncertainty characteristics in the normal vs. abnormal (heat based) conditions. The figures denote the estimated entropy states according to the time series between normal (5:00 and 17:00), vs. abnormal (5:00 and 17:00). N = normal, Ab = abnormal in terms of heat_based experimental design. Note: for this realization, participant 2’s data was used which could be represented as the most similar scores with the arbitrarily normalized scores from the averaged all participants’ scores as a function of the frequency competition [*SDϕ* (rad)].

**Figure 4.**
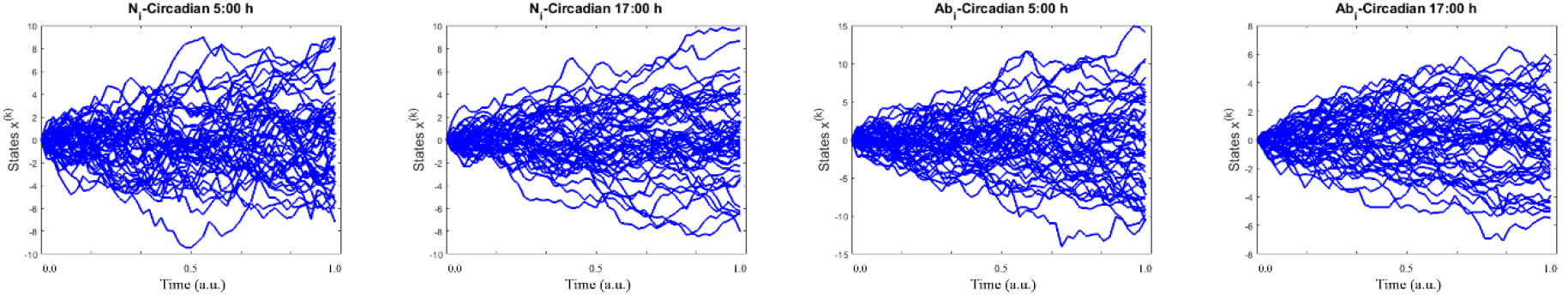
Uncertainty characteristics in the normal vs. abnormal (ice based) conditions. The figures denote the estimated entropy states according to the time series between normal (5:00 and 17:00), vs. abnormal (5:00 and 17:00). N = normal, Ab = abnormal in terms of ice_based experimental design. Note: for this realization, participant 5’s data was used which could be represented as the most similar scores with the arbitrarily normalized scores from the averaged all participants’ scores as a function of the frequency competition [*SD_ϕ_* (rad)].

The result shows the biological stability depending on the circadian time point, including the temperature [artificially perturbed body core temperature caused by the heated (iced) vest] perturbation. As shown in this figure 3, the main effect in the heated temperature perturbations was [*F*(1, 3) = 1.301, *η*^2^ = 0.961, (*p* < 0.258)]. The main circadian effect was [*F*(1, 3) = 20.531, *η*^2^= 15.166, (*p* < 0.001)], and the significant temperature perturbation by the circadian cycle on the biological motor synchrony disorder was [*F*(1, 3) = 3.453, *η*^2^= 2.551, (*p* < 0.068)]. As shown in the figure 4, the main effect in the iced temperature perturbations was [*F*(1, 3) = 1.211, *η*^2^= 0.861, (*p* < 0.275)]. The circadian main effect was [*F*(1, 3) = 23.041, *η*^2^= 43.317, (*p* < 0.001)], and the significant temperature perturbation by the circadian cycle on the biological motor synchrony disorder was [*F*(1, 3) = 4.264, *η*^2^= 3.035, (*p* < 0.043)]. These results indicate that although the participants exhibited significantly greater levels of entropy in 5:00 a.m. conditions compared to the 17:00 p.m. conditions, in both the normal and the abnormal conditions (circadian effect), the temperature-associated disorder difference between a.m. and p.m. was intensified during artificially increased body core temperature (interaction effect).

Estimated dynamics from the relative phase between the two limbs, oscillatory bimanual coordination was affected by the temporal locus during the circadian cycle. Results at this biological scale correspond to a theoretical study which has shown that the rate of entropy production is changed when a new energy source is accessed via a nonequilibrium phase transition process (Frank, 2011).

### Remarks of the results (theoretical implications 1)

The biology may be a complex sensor of its environment that can very effectively adapt itself in a broad range of different circumstances. Organism may convert internal energy of itself so efficiently that if it were to produce what is physically (thermodynamically) possible (England, 2013). These results from Experiment 1, 2, and 3 reflect that accessing a new energy source differs as a function of the circadian cycle, access that can be manipulated by a temporary thermal manipulation. A generalization of this model may not distinguish organism (x) and environment (y), but points to relevant characteristics that may include the context or circumstances in which physical dynamics takes place as a mutually constraining factor. Because, while (a) circadian effect divided by the artificially perturbed temperature manipulation is not constant according to different day-night temperature cycle, (b) circadian effect divided by the ordinary temperature seems may be constant according to the different day-night temperature cycle.

## Part 2: Systemic Proof of the Context-Dependency

Pervasive interconnectedness—everything is connected with some other thing or things (Mahner & Bunge, 1997) –suggests that behavior is adapted to perceiving both the nested environmental properties and one’s own nested behaviors, a union that organizes actions on surrounding circumstances (Reed, 1996). The discovery of direct and robust relation between biological aspects (body temperature and motor synchrony) an environmental process (circadian temperature cycle) may echo adaptation of our biological system to the environment.

These relations of both inner (bi-manual coordination) and outer (circadian temperature cycles) sources will provide a sign post with address intelligence as a natural consequence of the principles pertaining to the emergence of functional systems. Thus, physical principles might be methodologically reducible to any level of things (Iberall, 1977), and in this sense, it will motivate methodological and measurement advances which today constitute a strategy for examining self-organization in biological systems. The effect must extend the reach of thermodynamic theory and perhaps identify general principles that apply more broadly to complex systems.

### Conditioning main variables with regards to the components’ relationships

A core cycles of our system were influenced by temperature embedding on 24 hour light-dark oscillation (called circadian rhythm) according to the observation (Part 1); biochemical, physiological, or behavioral processes that persist under constant conditions with a period length of ~24 hours (Maury, Ramsey, Bass, 2010). We think that it is worth taking engaged variables when it comes to the experimental measurement and defined relatively independent processes: “circadian” and “temperature.” Simultaneously, these patterns or characteristics were constrained by the dynamical criteria of stability pattern when it comes to such systems possess a static equilibrium point, which can be called a point attractor (Kugler, Kelso, & Turvey, 1980).

As discussed above definition about the *ϕ* (state of the coupling strength given time), we say that these two objects are entangled which means knowing something about one causes we to know something about the other. It means the source of a relative phase is energetically emitting two correlated states in opposite directions. Suppose we take a particle (*φ*) in the state 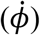 and subjected to experiment with two possible outcome (different directions) states.

The emission of one state is followed quickly by the second, and they share the same plane of stability, one up and one down. The measurement calls these parameters (and label the outcomes) as the biological coupling strength 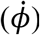, positive (+ = 1) and negative (− = 0). It assumed that a pair of coupling one-half states formed somehow in the singlet coupling state, moving freely in opposite directions. As it is, in classical mechanics, if the measurement of the component is one vector (>), the value of one side must yield the value of the other, opposite, side.

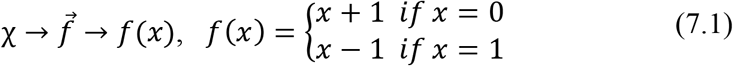

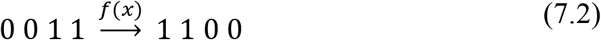

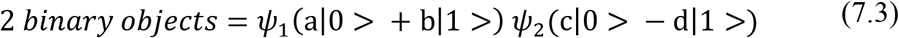

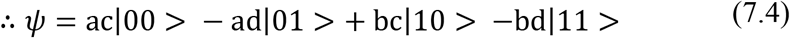

This must be state 0 plus 1 for the first particle (*ψ*_1_), with the second particle (*ψ*_2_) being in state 0 minus 1. This is the term 0, 0 for ac, plus the term 0,1 for ad with a minus ad, plus the term 1,0 for bc, and finally plus the term 1,1 for bd.

Because this logic can predict in advance the result of measuring any chosen component of 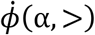 and 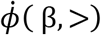, by previously measuring the same component of 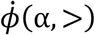 and 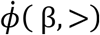, it follows that the result of any such measurement must actually be predetermined. The interpretation devises mechanical correlations such that each vector variable 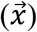 is a single continuous parameter, but the measurements of these parameters made in state (Ω) have no influence on those in other states. The expected value of the components is then determined as follows;

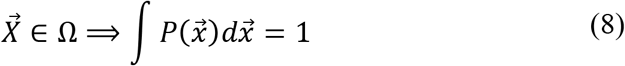

Here, 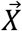 equals the parameter in the state space (Ω). 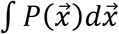 is the normalized distribution of these parameters, stating that the system is in a certain interval (*η,η* + *dη*). However, the present experimental design has another vector apart from the “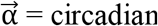” and “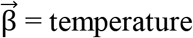” factors, which are always embedded in the system’s state as an interconnected component known as “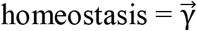.” According to the circadian temperature correlates of performance, a circadian temperature component and a homeostatic process are assumed to be mutually interdependent of both processes (Carrier & Monk, 2000). They are relatively independent but always correlated each other in that they are always communicating with one another regardless of how far apart they are according to the same processes. As it is, with normalization 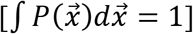 mapping onto the three components of β, α, and the value of γ, with the measurement variable 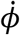, the results found different component conditions of biological states (entropy) have significant context dependency on each variable combination (see the results of the Part 1).

To describe the collections of the three macroscopic processes results, the interpretation of this finding undertakes a putative approximation. If a measurement considers the symmetric coupling strength 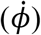 as positive (+) and negative (-) with a value of 1 and 0 (to simplify a two-by-two matrix), respectively, for the state of the particle, the measurement can reach a possible outcome function having three components (sine waves during the day).

Thus, the following binary objects function as a unit vector (*λ*), meaning that basic logic of this measurement takes the state of 0, resulting in 1. On the other hand, it takes 1 and gives 0.

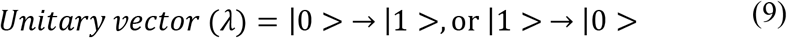

Considering the possible outcomes of these components underlying the unitary vector operation, there must be eight possibilities (see Table 7; parameter). Let this measurement considers a situation where *ϕ* faces the process of the first component with a change but faces the next process with no change, labelling this as 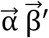(the circadian changes and not the temperature); that is, α equals 1 and β equals 0. There are two rows in a table for which α equals 1 and β equals 0. Next, if considering the situation where *ϕ* passes process 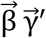 (a temperature changes and no homeostasis), this logic has two rows where β equals 1 and γ equals 0. Finally, consider the passing of 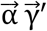 (a circadian change and no homeostasis), the logic also has two rows where α equals 1 and γ equals 0.

**Table 4:** Data collection for the experiment 3. 2 conditions, 8participants, 2 circadian points, 6 trials at each circadian point. Note: Participants were asked to swing in-phase of their limbs in different anatomy points [192 data set (3-level: wrist, elbow, and shoulder)], but used only wrist joint data (64 set) for analysis. Duration of each trial is 1 minute and 5 minutes rest interval between trials.

| Condition | Participants (N) | Circadian points | Trials at each circadian | Task/rest (min) |
| --- | --- | --- | --- | --- |
| Normal | 8 | 5:00 | 6 | 1/5 |
|  |  | 17:00 |  |  |
| Abnormal<br>(ice based) | 8 | 5:00 | 6 | 1/5 |
|  |  | 17:00 |  |  |

**Table 5:** Averaged entropy production from normal and abnormal (heat-based) day-night temperature effects. *N*(*I*) = number of case indexed by the calculation of (***w*_1_** + ***w*_2_** / 2), *AVER* = averaged entropy production; *STDEV* = averaged variability from the entropy production; *SES* = standard error score.

|  | Circadian N_5:00 | Circadian N_17:00 | Circadian Ab_5:00 | Circadian Ab_17:00 |
| --- | --- | --- | --- | --- |
| $N(I)$ | 8 | 8 | 8 | 8 |
| $AVER(H)$ | 0.410 | -0.165 | 0.564 | -0.809 |
| $STDEV(H)$ | 0.651 | 0.664 | 0.627 | 0.745 |
| $SES$ | 0.230 | 0.235 | 0.222 | 0.264 |

**Table 6:** Averaged entropy production from normal and abnormal (ice based) day–night temperature effects. *N*(*I*) = number of case indexed by the calculation of (***w*_1_** + ***w*_2_** / 2), *AVER* = averaged entropy production; *STDEV* = averaged variability from the entropy production; *SES* = standard error score.

|  | Circadian N_5:00 | Circadian N_17:00 | Circadian Ab_5:00 | Circadian Ab_17:00 |
| --- | --- | --- | --- | --- |
| $N(I)$ | 8 | 8 | 8 | 8 |
| $AVER(H)$ | 0.404 | -0.172 | 0.608 | -0.840 |
| $STDEV(H)$ | 0.446 | 1.031 | 0.518 | 0.993 |
| $SES$ | 0.158 | 0.365 | 0.183 | 0.351 |

**Table 7:** Visualizing the number of objects with combination outcomes. Notice: as indicated by these lines, whenever 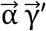 occurs either 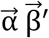 or 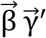 must also happen, conversely, table for which 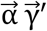 happens but either 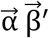 or 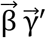 does not happen.

|  |  | 1 | 2 | 3 | 4 | 5 | 6 | 7 | 8 |
| --- | --- | --- | --- | --- | --- | --- | --- | --- | --- |
| Parameters | $\vec{\alpha} = \text{Circadian}$ | 0 | 0 | 0 | 0 | 1 | 1 | 1 | 1 |
| | $\vec{\beta} = \text{Temperature}$ | 0 | 0 | 1 | 1 | 0 | 0 | 1 | 1 |
| | $\vec{\gamma} = \text{Homeostasis}$ | 0 | 1 | 0 | 1 | 0 | 1 | 0 | 1 |
| Conditions | $\vec{\alpha} \vec{\beta}'$ | | | | | ✓ | ✓ | | |
| | $\vec{\beta} \vec{\gamma}'$ | | | ✓ | | | | ✓ | |
| | $\vec{\alpha} \vec{\gamma}'$ | | | | | ✓ | | ✓ | |

**Table 8:** Representing the process (first, second, and third) with properties. α = circadian process. β = temperature process. γ = homeostatic process. (↑), (↓) object’s (ϕ) state up or down based on *g*(*x*) function.

| Angle \ Process | $0_\phi$<br>$g(x)_\alpha$ | $\theta_\phi$<br>$g(x)_\beta$ | $2\theta_\phi$<br>$g(x)_\gamma$ |
| --- | --- | --- | --- |
| First | (↑) | (↓) | |
| Second | | (↑) | (↓) |
| Third | (↑) | | (↓) |

### Proving components’ relationship by way of inequality

Assuming these combinations, the interpretation can write the formula considering the probability distribution of the three unit vectors, as follows;

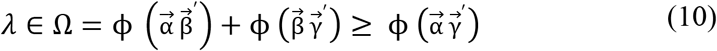

Because the statistical predictions of this correlation cannot be arbitrarily equal to the vectors combinations (Bell, 1964), the formal proof of this can be surmised as shown below.

Figure 8 shows some of the interconnected relationship of the objects. The parameters α, β, and γ refer to measurements of the 1:1 frequency-locked synchronization phase 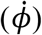. If any property enters another property first, the component can subsequently include the others and change simultaneously. The measurement has calculations which keep track of the area of the circles 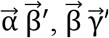, and 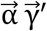. These relationships can be indicated as the given proportion (A; circle area).

**Figure 5.**
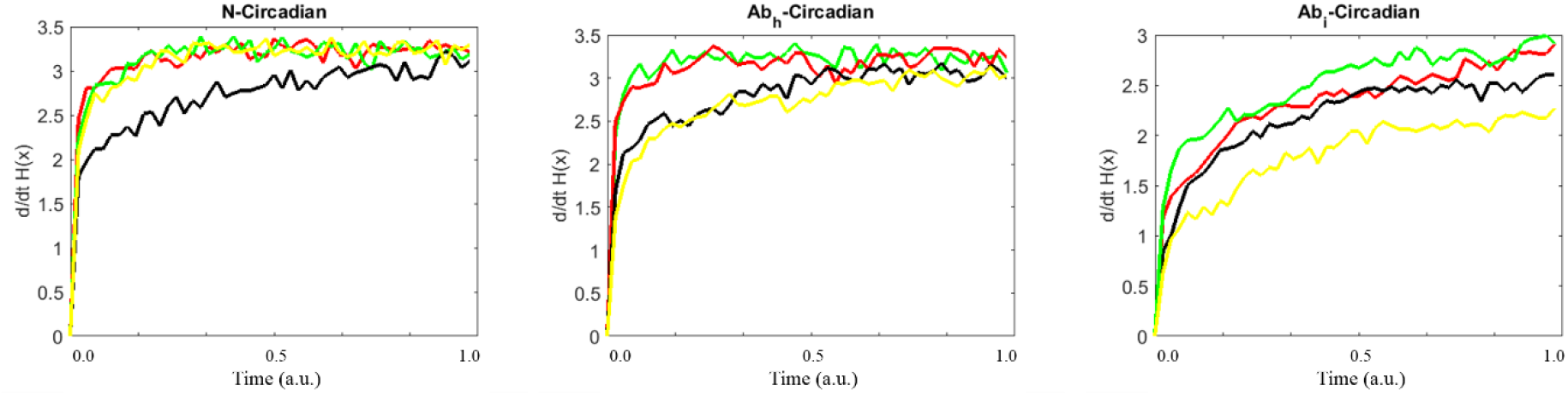
Entropy maximizations for the all uncertainty characteristics. Each figure denotes the estimated entropy forces according to the time series for each experimental design. The figure on the left denotes the entropy forces of the normal condition (red line = 5:00, green line = 12:00, black line = 17:00), which data was from the Figure 2. The figure in the middle denotes the entropy forces of the heat-based normal (red line = 5:00, black line = 17:00) vs. abnormal (green line = 5:00, yellow line = 17:00) conditions, which data was from the Figure 3. The figure on the right denotes the entropy forces of the ice-based normal (red line = 5:00, black line = 17:00) vs. abnormal (green line = 5:00, yellow line = 17:00) conditions, which data was from the Figure 4.

**Figure 6.**
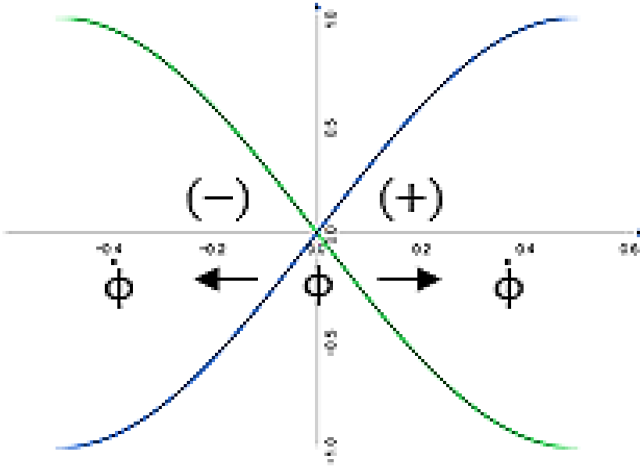
Schematic drawing of the measurement system. x_axis means time and y_axis represents space of the two entangled particles (*ϕ*). The particle departs from a source and move apart in opposite directions (±). It is detected by a correlated two space with two possible outcome states 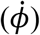.

**Figure 7.**
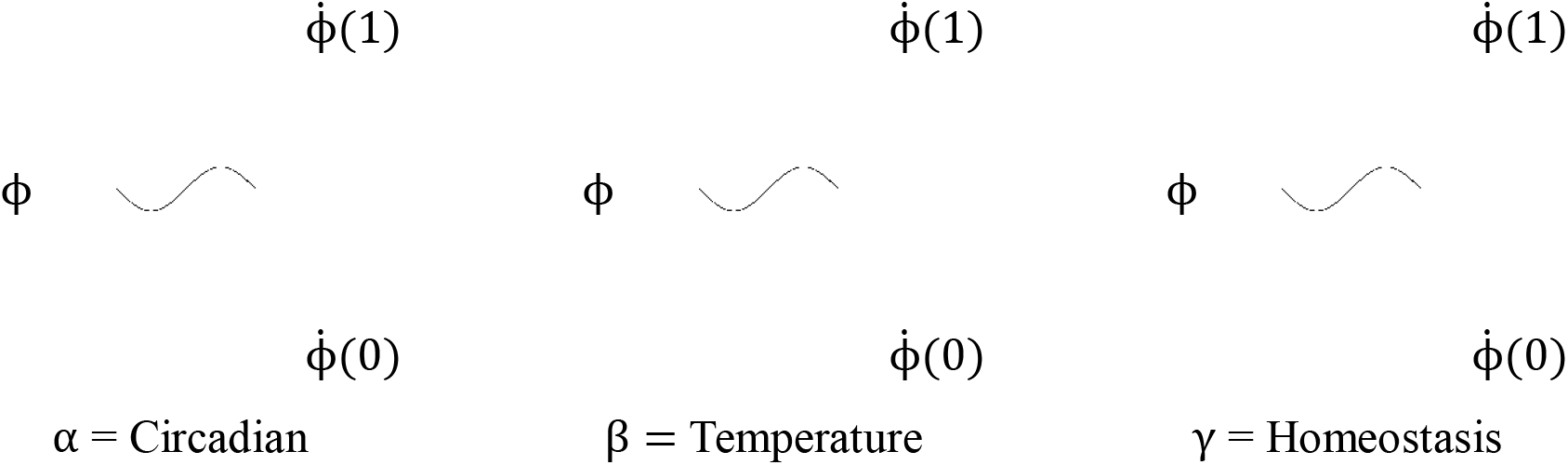
Illustration of a three double-valued parameters specification of a physical property. α means circadian component. β denotes temperature component. γ represents homeostatic component. 1 is outcome of the 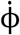 change, while 0 is not change.

**Figure 8.**
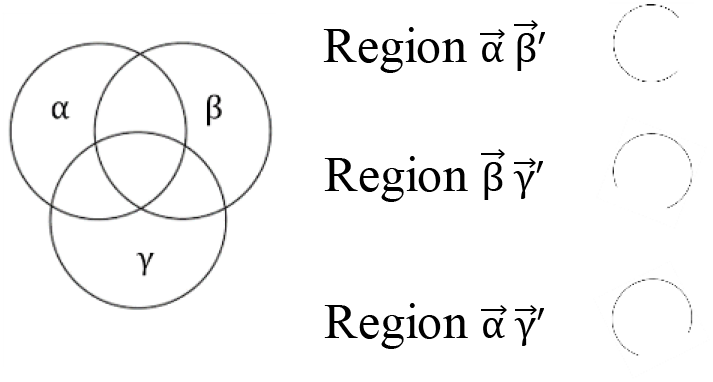
Representing the number of objects with properties. α = circadian component. β = temperature component. γ = homeostatic component.

The area of the circle of the 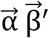 proportion plus the area of the circle of the 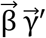 proportion must be greater or equal to the area of the circle of the 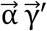 proportion. As indicated by the combination table, this is because the last combination 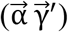 cannot occur without one and the other two combinations 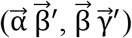 also transpiring. However, it is possible for one of the first two combinations 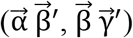 to occur without the third combination 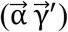 transpiring. Their two frequencies (rows) in the table 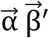 or 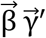 are captured, while 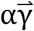 is not. Therefore, the measurement has an inequality on the left side 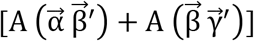 which is greater than or equal to that on the right side 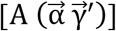.

Because this assumption is true for any collection of relationships, this logic assumes a type of realism in the proof that these parameters have properties regardless of whether they are measured or specified or not; the logic rewrites these proportions in terms of the sector area (*Â*) of each circle.

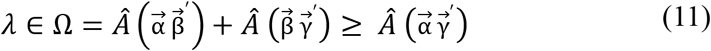

Based on the above logic, this representation simulates the sector area from each circle for which two lines have identical radiuses. These two radiuses from the center of the circle create the area of the circle. It is necessary to measure the proportion of the large area or the sector of the circle because, as shown in the diagram (Figure 9), the proportion of the 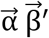 area is indicated as the large part of the circle. Next, the representation gives the radius tentatively as a value of two inches and give the large area of the circle as ¾ (the small area is ¼). The area of the circle is pi (π) multiplied by the radius squared (*A* = *πr*^2^). Given that the overall angle of the circle α is 360 degrees, the 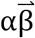 portion of the circle as referred to here is 270 degrees (small area: 90 degree). If the measurement takes this fraction and multiplies it by the area of the entire circle, the result is as follows;

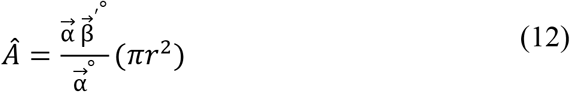

**Figure 9.**
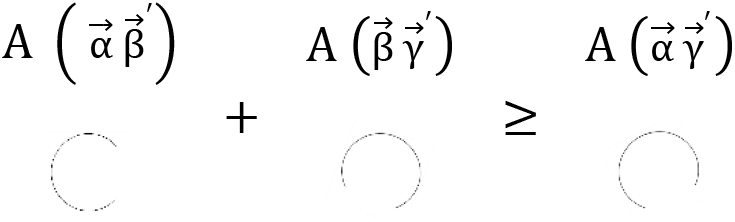
Mapping of the objects with the area of the circles. α = circadian component. β = temperature component. γ = homeostatic component.

**Figure 10.**
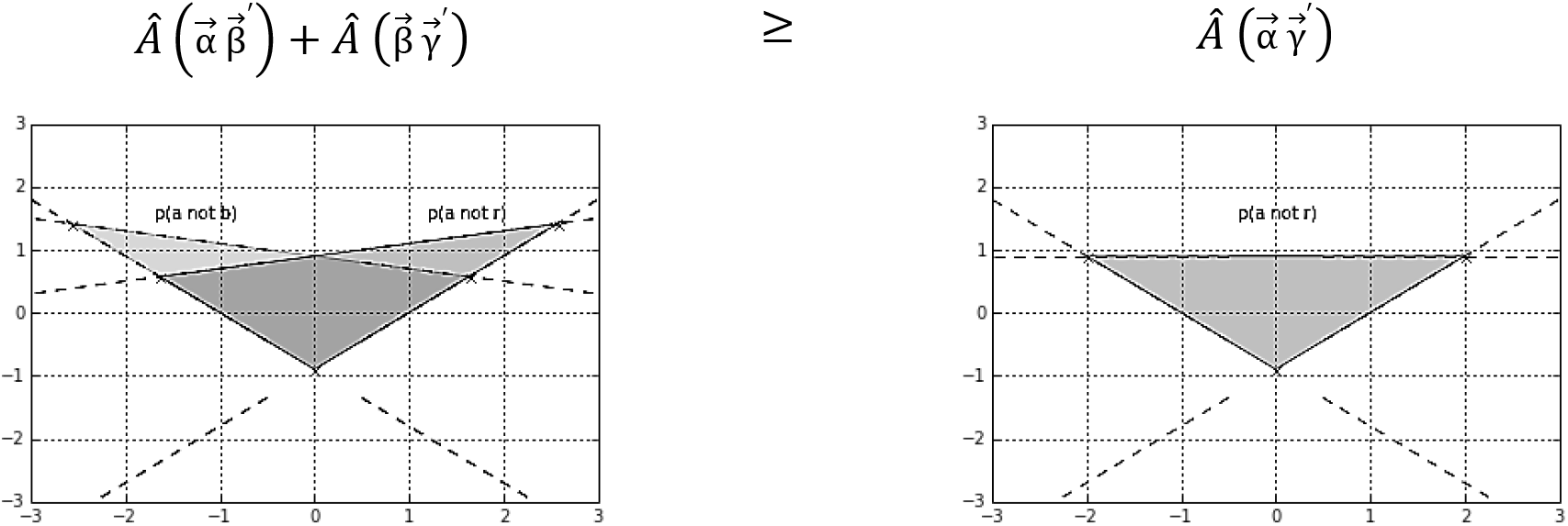
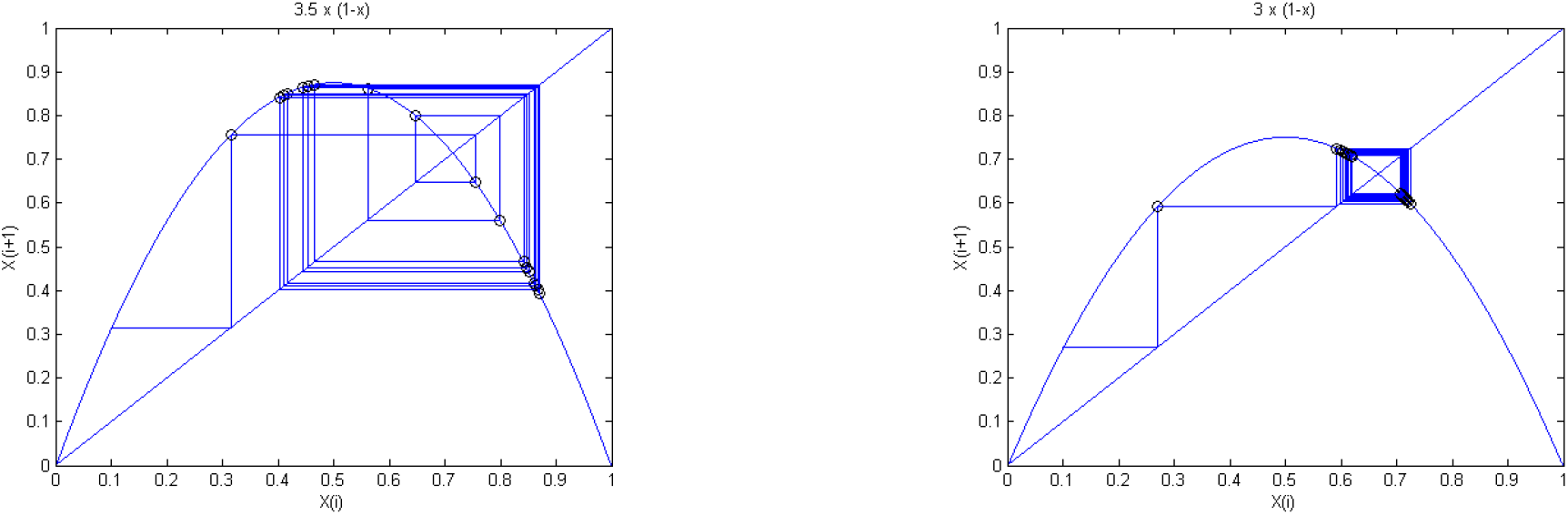
Representing the probability of objects with Cartesian coordinate system. Left side of the upper plot = combined proportion of 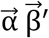 and 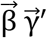; [relative portion: 3.5/4(*π*4) = 10. 99 sq.in.]. Right side of the upper plot = proportion of 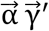; [relative portion: 3/4(π4) = 9. 42 sq.in.]. Bottom shows their inequalities of the system consideration of the complex behavior [Initial value of x is 0.1, iteration frequency 20, and *X*_*n*+1_ is *r_χ_n__*(1 – *X_n_*)]. Left plot is showing when parameter value of *r* equals 3.5 (combined proportion of 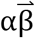 and 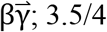), while right is showing the *r* is 3 (proportion of 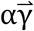 3/4).

**Figure 11.**
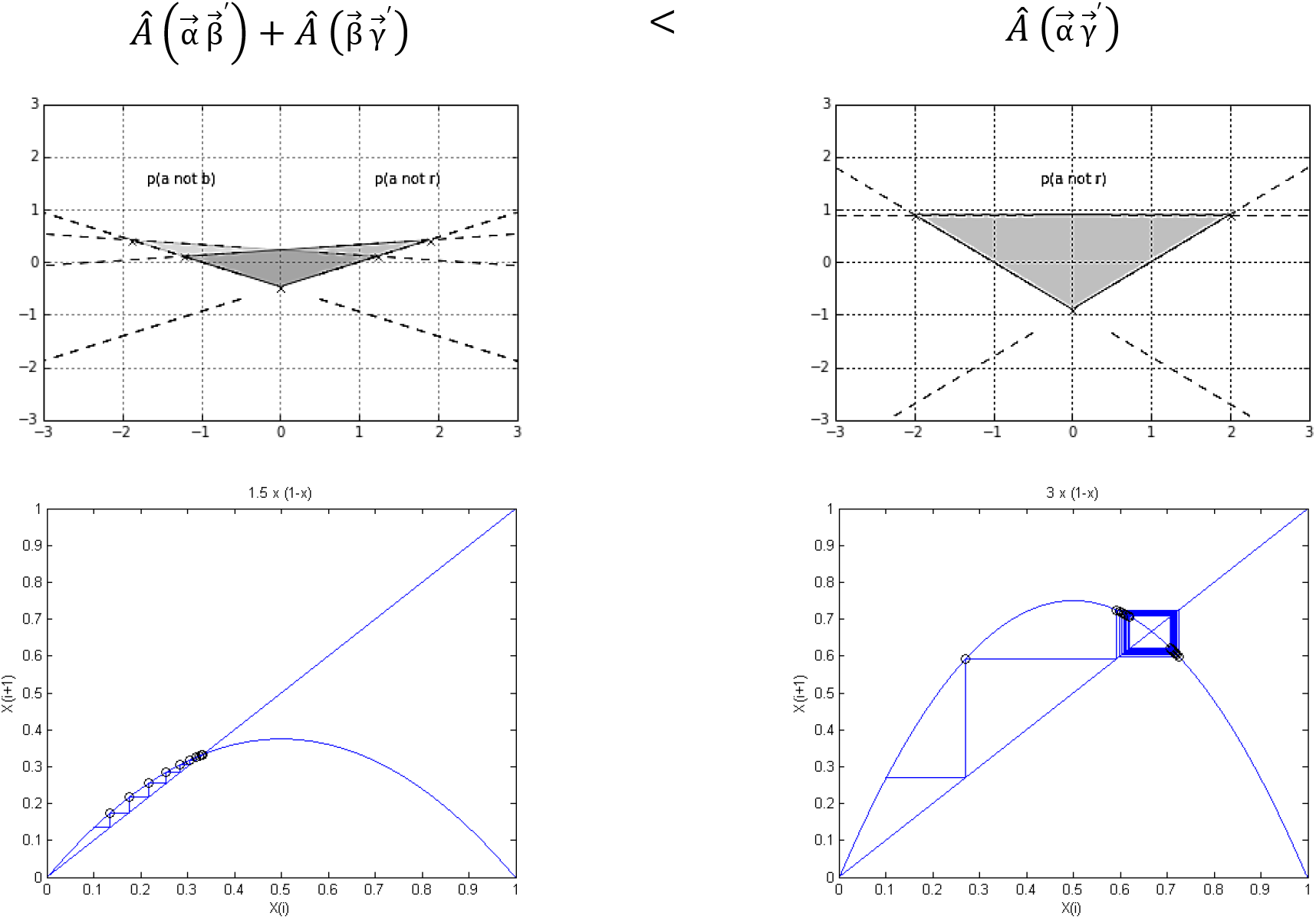
Representing the probability of objects with Cartesian coordinate system(Upper). Upper left = combined proportion of 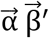 and 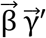; [relative portion: 1.5/4(*π*4) = 4. 71 sq.in.]. Upper right = proportion of 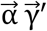; [relative portion: 3/4(*π*4) = 9. 42 sq.in.]. Bottom plots

Here, *Â* denotes the sector area of component α. 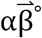 is a large portion of *θ*°(270 degree), and α° is the entire portion of *θ*°(360 degree). The equation for 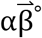 over α° requires the multiplication of the area of the entire circle, which is (*πr*^2^). If the measurement simplifies both the numerator and denominator of this fraction, the result is ¾, after which the logic multiplies this value by *π* multiplied by the radius (2) squared, giving a solution to the proportions of the large sector area of these three components [3/4(*π*4) = 9. 42 sq.in.]. These complex portions are represented via the Cartesian system for a linear relationship (complex system for a nonlinear), as shown below;

The fact that all objects each have all properties including the others corresponds to the changes. The main principle of this equation is the simultaneous measure, for instance, property 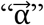 and property 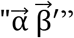 of the same property. These interpretations are taken as proof that our experimental feature by which the principles of the state of action and the antecedence do apply to the system. Regardless of how many trials are run, because we have three parameters, it will always be a case where left combinations (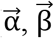, and 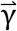) must greater than or equal to right combinations (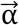 and 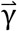), such as our perturbed morning entropy being greater than normal condition.

However, here is the experimental complex which must be proved regarding the temperature perturbed pm condition: the measurement assumes the experimental set again with wave function [*g*(*x*)] similar to when object (ϕ) faces the parameters with the states of their angles up (↑) or down (↓).

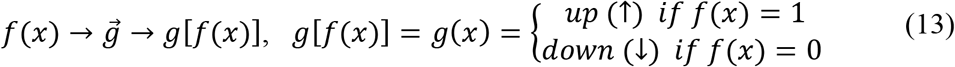

The first parameter 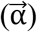 changes the state to up or down at 0, and 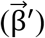 changes the state up or down at *θ*. The second parameter 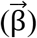 then changes the state up or down at 0, while 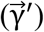 changes the state up and down at 2*θ*. The third parameter 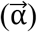 changes the state up or down at 0, and 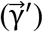 changes the state up and down at 2*θ*.

This function can simplify the inequality considering the experimental relative phase change at each parameter set as follows;

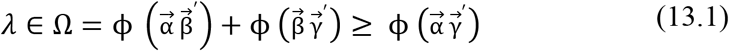

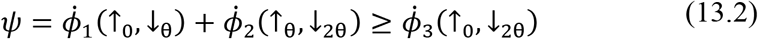

The expressions above assume that with 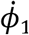, if the measurement of one side (the entangled relative phase) is up or down at 0, the opposite side of the state must be up or down at θ because they are entangled with a unitary vector operator. For 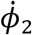, the next state of one side is then going to be up or down at θ, and the opposite side of the state must then be up or down at 2θ. For 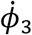, one side is up or down at 0, and the state of the opposite side must then be up or down at 2θ. This inequality can be computed with the common angle scales shown below,

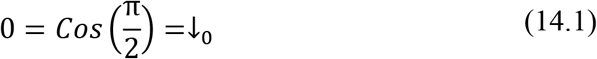

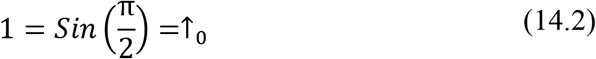

When the measurement refers to angles, the two common “scales” are degrees and radians. In degrees, the measurement uses the π notation. Understand that π (radians) is equivalent to 180 degrees, meaning that π/2 equals 90 degrees. In this case, the sine of π/2 equals 1/1, and when the sine of π/2 is 1, cos π/2 becomes 0. However, to prove this, because this logic cannot be defined by the relative phase calculation [*ϕ* = Δ*ω*(*t*) + *ϕ*(0)] simply as a π value (*ϕ* ≠ π), the change in the relative phase angle is replaced by a putative degree of theta (*θ*) after which each state is expressed as follows;

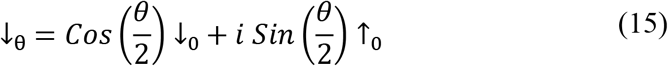

In this equation, if the phase angle is down at theta 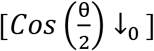, the probability that the phase being up at 0 is 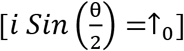. It should be ↓_θ_ because 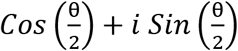 equals 1 (*i* is the imaginary unit). Thus, the probability of the first state phase being up or down at 0 is 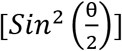. Likewise, the next probability that the phase is up or down at theta (↑_θ_) and up or down at 2 theta (↓_2θ_) must be identical, as the rotation of theta gives us an identical value.

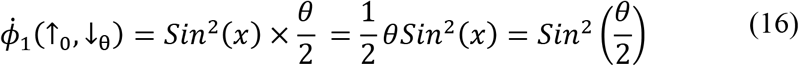

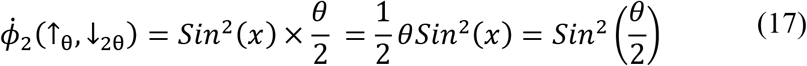

However, the probability that the phase is up at 0 (↑_0_) and down at 2 theta (↓_2θ_) is the sine squared of theta, as this state only takes a factor of 2 for theta.

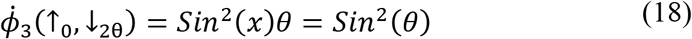

Based on these wave functions, the terms can be redefined as follows;

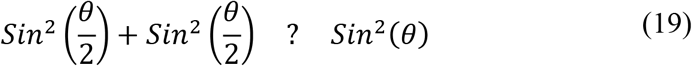

Hence, the sine squared theta is less than 1 (θ < 1), and this function can be simplified on the left side as follows;

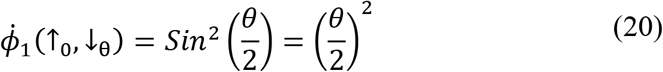

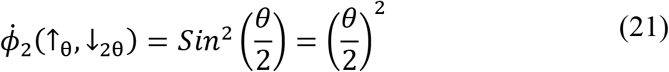

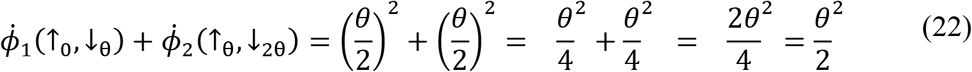

This can also be done on the right side, as follows;

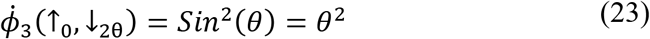

The measurement can finally be written in the terms below, to reevaluate the inequality.

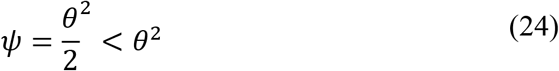

As shown above, the left-hand side 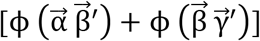 cannot be greater than or equal to the right-hand side 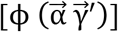.

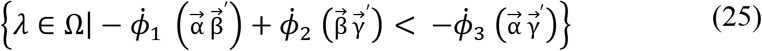

Using these observations, the measurement finds an explicit violation of a previous inequality (Harrison, 1982) and suggest this as empirical proof underlying the probability of temperature-perturbed pm events. This set solution can also be represented by the Cartesian system for linear relationships (complex system for a nonlinear), as follows; illustrate the complex behavior [Initial value of x is 0.l, iteration frequency 20, and *X*_*n*+1_ is *r_χ_n__*(1 – *X_n_*)]. Left plot is showing when parameter value of *r* equals 1.5 (combined proportion of 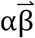 and 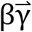; 1.5/4), while right is showing the *r* is 3 (proportion of 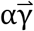; 3/4).

The result shows that the predictions of outcomes for the states by the system are inconsistent (Hensen, Bernien, Dréau, Reiserer, Kalb, Blok, … & Amaya, 2015; Wiseman, 2015) with the parameters (different combinations) and measurements (different calculations). In the context of interaction between all parameters and measurements, the deviation of phase inequality asserts a fundamental limit on the precision of certain pairs of physical position and momentum properties simultaneously. The remarkable possibility, this interpretation can estimate, is that whereas the relationship between objects’ energy and reaction rates obtains only under certain constraints, having derived inequality here indicates that the entangled objects’ states applies in cases where many degrees of freedom in the system states out of thermal equilibrium (England, 2013). These entangled features prove the heterogeneity of the system in an ecologically dependent process context.

### Remarks of the results (theoretical implications 2)

As evident from the related biological scale, we encourage the belief that there is no common prediction of bias for uncertainty toward the system of ecology. The space between antecedents and consequences is the heart of the process, and we must look at how the variables change. It is recursive in that each variable and process can affect each other depending on where in the flow of behavior one begins. A few scholars in science have investigated experimental models that guide the circadian process. Differential time-of-day variations for different tasks were observed under a normal day-night condition. No attempt was made to distinguish variations in performance due to endogenous circadian factors relative to those linked to the amount of time since awaking. Perhaps the main conclusion to be drawn from studies of the effects of the time of day on performance is that the best time to perform a particular task depends on the nature of that task (Folkard, 1983).

One study showed an early morning peak of mental arithmetic performance in children (Rutenfranz & Helbruegge 1957), while another study found an evening peak for this type of performance in highly practiced young adults (Blake, 1967). With a low working memory load, performance was correlated positively with the circadian rhythm of the body temperature (Folkard, Knauth, Monk, & Rutenfranz, 1967). The majority of the performance-related components (flexibility, muscle strength, short-term memory) appear to vary with the time of day. In particular, contemporary models of subjective alertness and performance efficiency view these variables as being determined both by a homeostatic process (number of hours since waking) and by input from the circadian timing system (Monk, Weitzman, Fookson, Moline, Kronauer, & Gander, 1983; Johnson et al., 1992). There is still much work to do before one can understand which performance tasks will show different time-of-day effects and which will define the mechanisms that underlie these differences.

Frequency locked 1:1 coupling embedded on a bi-manual pendulum 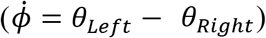 and the homeostatic process (temperature) embedded on the circadian 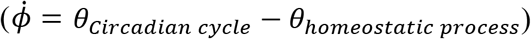 can be illuminated by a similar relative phase dynamic function. The result of the part 1 shows that the system’s stability decreases more at the circadian AM time when the system is perturbed as compared to the normal circadian AM condition. However, the stability of the system increases more at the circadian PM time in spite of being perturbed as compared to normal circadian PM condition. What relations must hold for the formulae to be an interpretation of a biological state with respect to an environmental process in the system? Assumption of this interpretation does not intend to have any precise, predicative knowledge corresponding to the systemic proof in part 2. For a biological state to be related to the environmental process represented by the system, the system must be considered in conjunction with its basis. This is simply the relationship between one and another with infinite distinct representations of the system’s productivities.

## Conclusion

Inquiry into the possibility of relating perception-action to dynamics began in the 1970s with the problem of coordination: could a principled dynamical account be given of fundamental rhythmic capabilities involving multiple joints, scores of muscles, and millions of cells? Efforts to address this question invoked the concepts and tools of nonlinear dynamics (e.g., Kugler & Turvey, 1987). One useful approach to the relationship between dynamics and self-organization is provided by *Homeokinetics* (e.g., Soodak & Iberall, 1978), a strategy that looks for cycles at all time scales and aims to show how interacting cyclic processes account for the emergence of new entities, many of which are similarly cyclic. The central idea is that the cycles have interacted to create self-replicating living systems abiding particular cyclicities (Iberall, 2001). Attunement to these cycles is integral to life. Circadian rhythms are cycles of particular prominence in contemporary research on living things. Rather than crediting the rhythms to “clock genes,” the dynamic approach considers them as an emergent property of the system.

They are found in most living things – especially in humans. These rhythms are not to be confused with the biological system. However, they are related in that the circadian system drives the biological system. The circadian rhythms that control biological rhythms are grouped interacting molecules in cells throughout the body that act in sync with each other. This has been being a major issue, and it is hugely influential on us because we are in effect physical, emotional and performance systems in our bodies (Monk et al., 1983). Contemporary models of subjective alertness and performance efficiency view these variables as being determined both by a homeostatic process (amount of hours since waking) and by input from the circadian timing system (Johnson et al., 1992). However, there is still much work to do before one can understand which performance tasks will show different time-of-day effects with various environmental variables and which mechanisms underlie these differences. Corresponding to a theoretical study about the rate of the stability of a system and how it can vary when a new energy source is accessed via a nonequilibrium phase transition process (Frank, 2011), the results from this experiment appear to reconfirm that access differs as a function of thermodynamic variables (circadian temperature), which means access that can be manipulated by temporary thermal manipulation methods.

One must know the larger system to characterize the smaller system, but one cannot know the larger system in the absence of the characterization of the smaller system. Such that may be called this the problem of impoverished entailment (Shaw, 2001). This does not address directly, however, the question as to why the baseline of the different behavioral modes differ as they do. The ultimate significance of a recurrence of behavioral modes is that it allows the animal to balance the entropic (order-reducing) degradations associated with the processes, thereby ensuring persistence of its characteristic forms and functions (Iberall & Soodak, 1987). The biology may be a complex sensor of its environment that can very effectively adapt itself in a broad range of different circumstances. Organism may convert internal energy of itself so efficiently that if it were to produce what is thermodynamically possible (England, 2013). As it is, generalization may not distinguish organism (x) from environment (y), or larger (biology) factors from smaller (anatomy); such systems may include the context or contexts in which physical dynamics can exist as a mutual factor. This suggestion would not response the question of where or how the system emergent, but provides a coherent account between biology and environment.

Everything is connected with some other thing or things (Mahner & Bunge, 1997), and adapt their behavior perceiving both the nested environmental properties and their own nested behaviors as a union (Turvey et al., 1981) in order to organize actions according to their surrounded circumstance (Cariani, 1993; Ford, 2008; Reed, 1996; Strong & Ray, 1975). Which may reflect that the mechanism of the system’s state that is not a specific component of special properties but a general co-activity encompassing all components (Gottlieb, 2000). It will be a reduction of physical principles (Iberall, 1977) classifying them as having a nervous system is not necessary condition. Spontaneously evolved engineering of this internal characteristics embedding in external context awaits future investigation so that we labeled these sorts of things to be deserved calling as physical intelligent (Turvey & Carello, 2012).

## Acknowledgments

This research was supported by an NSF Grant BCS-1344725 (INSPIRE Track 1), and the National Research Foundation of Korea (2016K2A9A1A02952017). All of them provided written informed consent to the study approved by the local ethics committee (SNUIRB No.1509/002-002) and conformed to the ethical standards of the 1964 Declaration of Helsinki (Collaborative Institutional Training Initiative Program, report ID 20481572).

